# Protein Interactomes Identify Distinct Pathways for *Streptococcus mutans* YidC1 and YidC2 Membrane Protein Insertases

**DOI:** 10.1101/2020.04.07.031013

**Authors:** Patricia Lara Vasquez, Surabhi Mishra, Senthil K. Kuppuswamy, Paula J. Crowley, L. Jeannine Brady

## Abstract

Virulence properties of cariogenic *Streptococcus mutans* depend on integral membrane proteins. Bacterial protein trafficking involves the co-translational signal recognition particle (SRP) pathway components Ffh and FtsY, the SecY translocon, and membrane-localized YidC chaperone/insertases. Unlike *Escherichia coli, S. mutans* survives loss of the SRP pathway. In addition, *S. mutans* has two *yidC* paralogs. The *ΔyidC2* phenotype largely parallels that of *Δffh* and *ΔftsY* while the *ΔyidC1* phenotype is less severe. This study defined YidC1 and YidC2 interactomes to identify their respective functions alone and in concert with the SRP, ribosome, and/or Sec translocon. A chemical cross-linking approach was employed, whereby whole cell lysates were treated with formaldehyde followed by Western blotting using anti-Ffh, FtsY, YidC1 or YidC2 antibodies and mass spectrometry (MS) analysis of gel-shifted bands. Cross-linked lysates from WT and *ΔyidC2* strains were also reacted with anti-YidC2 antibodies coupled to magnetic Dynabeads™, with co-captured proteins identified by MS. Additionally, C-terminal tails of YidC1 and YidC2 were engineered as glutathione-S-transferase fusion proteins and subjected to 2D Difference Gel Electrophoresis and MS analysis after being reacted with non-cross-linked lysates. Results indicate that YidC2 works in concert with the SRP-pathway, while YidC1 works in concert with the SecY translocon independently of the SRP. In addition, YidC1 and/or YidC2 can act alone in the insertion of a limited number of small integral membrane proteins. The YidC2-SRP and YidC1/SecY pathways appear to function as part of an integrated machinery that couples translation and transport with cell division, as well as transcription and DNA replication.

**Importance:** *Streptococcus mutans* is a prevalent oral pathogen and causative agent of tooth decay. Many proteins that enable this bacterium to thrive in its environmental niche, and cause disease, are embedded in its cytoplasmic membrane. The machinery that transports proteins into bacterial membranes differs between Gram-negative and Gram-positive organisms. One important difference is the presence of multiple YidC paralogs in Gram-positive bacteria. Characterization of a protein’s interactome can help define its physiological role. Herein, we characterized the interactomes of *S. mutans* YidC1 and YidC2. Results indicate that YidC1 and YidC2 have individualized functions in separate membrane insertion pathways, and suggest putative substrates of the respective pathways. Furthermore, *S. mutans* membrane transport proteins appear as part of a larger network of proteins involved in replication, transcription, translation, and cell division/cell shape. This information contributes to our understanding of protein transport in Gram-positive bacteria in general, and informs our understanding of *S. mutans* pathogenesis.

## Introduction

Dental caries, is the most common infectious disease in the world [1]. Tooth decay occurs when acidogenic bacteria on the tooth surface take up and ferment dietary sugars, producing organic acids that cause enamel demineralization. A major agent of caries, *Streptococcus mutans*, is acidogenic and aciduric enabling this species to tolerate acid end products and outcompete other oral microbiota. *S. mutans* displays inherent characteristics that promote dominance in its ecological niche, including efficient carbohydrate uptake and fermentation, sucrose-dependent and sucrose-independent adhesins leading to biofilm formation, robust acid tolerance mechanisms, and quorum-sensing systems involved in bacteriocin production and genetic competence [2]. These processes depend on integral membrane proteins, and/or membrane-associated proteins. *S. mutans’* competitive advantage and virulence attributes stem from its ability to sense and adapt to the harsh conditions it faces in the oral cavity. Efficient protein transport into and through the membrane is an essential aspect of this adaptability.

In bacteria, many integral membrane proteins are inserted into the cytoplasmic membrane co-translationally using the Signal Recognition Particle (SRP) pathway conserved in all living cells [reviewed in [3]]. The SRP binds hydrophobic signal sequences of nascent polypeptide substrates as they emerge from the ribosome. The bacterial ribosome-nascent-chain (RNC) complex is targeted to the membrane via a transient interaction of the SRP protein Ffh with the bacterial SRP receptor, FtsY. This docks the RNC with the SecYEG translocon pore, and enables translocation of the substrate into the membrane concomitant with translation. In addition to SecYEG, the integral membrane protein YidC also participates in membrane protein integration [4]. YidC belongs to the Oxa/Alb/YidC family of insertases found in mitochondria, chloroplasts, and bacteria. Membrane biogenesis has been most widely studied in the Gram-negative bacterium *Escherichia coli*; however, studies in Gram-positive bacteria such as *S. mutans* and *Bacillus spp.* have revealed differences in the translocation machineries of Gram-negative and Gram-positive organisms [5]. Importantly Gram-positive bacteria almost universally encode two, or occasionally more, YidC paralogs. Gram-negative organisms possess a single YidC.

The SRP pathway is dispensable in *S. mutans*, although its disruption results in growth impairment, environmental stress-sensitivity, and diminished genetic competence [6] [7]. Deletion of *S. mutans yidC2* causes a similar phenotype, whereas deletion of *yidC1* appears less detrimental [7-10]. YidC is essential in *E. coli* [11]. *S. mutans* survives elimination of *yidC1* or *yidC2*, but a double mutant is not viable. Nor is a double *yidC2*/*ffh* deletion mutant. In contrast, an *ffh/yidC1* mutant is viable, albeit severely stress sensitive and growth impaired [10]. These results suggest synthetic lethality and functional redundancies between the SRP pathway and YidC2, and between YidC1 and YidC2. While YidC1 and YidC2 apparently substitute for one another in some cases, distinct functional activities have been identified. YidC1 impacts cell surface biogenesis and bacterial adhesion more than YidC2, while YidC2 impacts cell wall biosynthesis and localization of penicillin binding proteins to the division septum [9, 10]. YidC1 and YidC2 demonstrate 27% amino acid homology and 48% similarity, each having six predicted transmembrane (TM) domains in the preprotein, and five in the mature insertase (TM2-TM6). The cytoplasmic C-terminal tail of YidC2 is longer and more highly charged than YidC1’s, and appending the YidC2 tail onto YidC1 enables the chimeric protein to partially complement *ΔyidC2* stress sensitivity [8]. Furthermore, gain of YidC2-like function point mutations have been reported within TM2 of *S. mutans* YidC1 [10], as well as within TM2 of SpoIIIJ, the YidC1 homolog of *Bacillus subtilis* [12]. Thus, while there is functional overlap between Gram-positive dual YidCs, paralog-specific features are recognized. It is of interest, therefore, to compare *S. mutans* YidC1 and YidC2-related interactomes to define respective roles of each paralog in the physiology of this pathogen, and understand the reason for dual YidCs in Gram-positive bacteria in general.

*E.coli* YidC can work independently [11, 13, 14], in collaboration with the Sec machinery [15-17], and in collaboration with SRP pathway components [17, 18]. *E.coli* membrane proteins inserted by YidC alone are relatively few, and generally contain only one or two TM domains [19-24]. Insertion of larger membrane proteins requires the Sec machinery and YidC [18, 25]. Respective substrates of integrated YidC/SecYEG and YidC/SecYEG/SRP pathways are largely unknown. Comparison of the membrane proteomes of *S. mutans* wildtype and mutant strains lacking *ffh, yidC1, yidC2, or ffh/yidC1* suggested that its SRP pathway works in concert with YidC1 or YidC2 specifically, or with no preference, to insert most membrane-localized substrates [10]. In a few instances only the SRP pathway, or only YidC1 or YidC2, appeared to be required [10]. Past studies of Gram-positive YidCs have used genetic approaches comparing phenotypic differences between wild-type and mutant strains [7, 8, 12], or cross-complementation in heterologous systems [26-28]. Solved crystal structures of bacterial YidCs [29-35] have also facilitated investigations of insertase interactions with other protein transport machinery components [17, 36-40]. Such studies employed *in vitro* or highly defined systems and provided information regarding particular protein-protein combinations. In contrast, in the current study we utilized an unbiased screening approach to evaluate similarities and differences between protein interactomes of *S. mutans* YidC1 and YidC2 within whole cell lysates. This led to identification of potential common, as well as YidC1- or YidC2-specific, substrates, identified interaction networks including proteins associated with translation as well as transcription and DNA replication, and revealed that YidC2 operates in concert with the SRP pathway while YidC1 operates in concert with the Sec machinery independently of SRP.

## Results and Discussion

### Identification of potential binding partners of *S. mutans* YidC1, YidC2, Ffh, and/or FtsY in whole cell lysates by formaldehyde cross-linking and Western blot gel shift

As a first step toward identifying putative binding partners and/or substrates of YidC1, YidC2, or the SRP pathway we utilized the cell penetrating cross-linking agent, formaldehyde. After cross-linking, whole cell lysates were prepared, separated by SDS-PAGE and potential regions of interest were identified by Western blot with anti-YidC1, YidC2, Ffh, and FtsY-specific antibodies (Fig. 1A). Bands corresponding to YidC1, YidC2, Ffh, and FtsY were readily identified in both cross-linked and non-cross-linked samples. Western blotting also revealed several regions of gel-shifted antibody reactivity in the formaldehyde cross-linked sample compared to the non-cross-linked sample (Fig.1A). Three distinct gel-shifted regions were excised from corresponding Coomassie Blue-stained SDS-polyacrylamide gels for mass-spectrometric (MS) analysis (Fig. 1B). These included a high molecular weight region (∼ 200-250 kDa) reactive with anti-YidC2, Ffh, and FtsY, but not anti-YidC1 antibodies, a middle region of ∼40-45 kDa reactive with anti-YidC1 and anti-YidC2 antibodies, and a lower region of ∼30-33 kDa reactive only with anti-YidC1 antibody (Fig. 1A). Proteins present in the upper, middle and lower MW gel slices of the cross-linked sample, but not the non-cross-linked sample, are summarized in Table S1. Initially, MS analysis was performed only on the upper and middle molecular weight regions, but because relatively few proteins were identified in that experiment, we moved on to the immunocapture approach described below to improve sensitivity. During that time period a more sensitive mass spectrometer became available and the experiment was repeated with analysis of all three molecular weight regions. The results presented represent the combined data from both experiments.

**Fig. 1.**
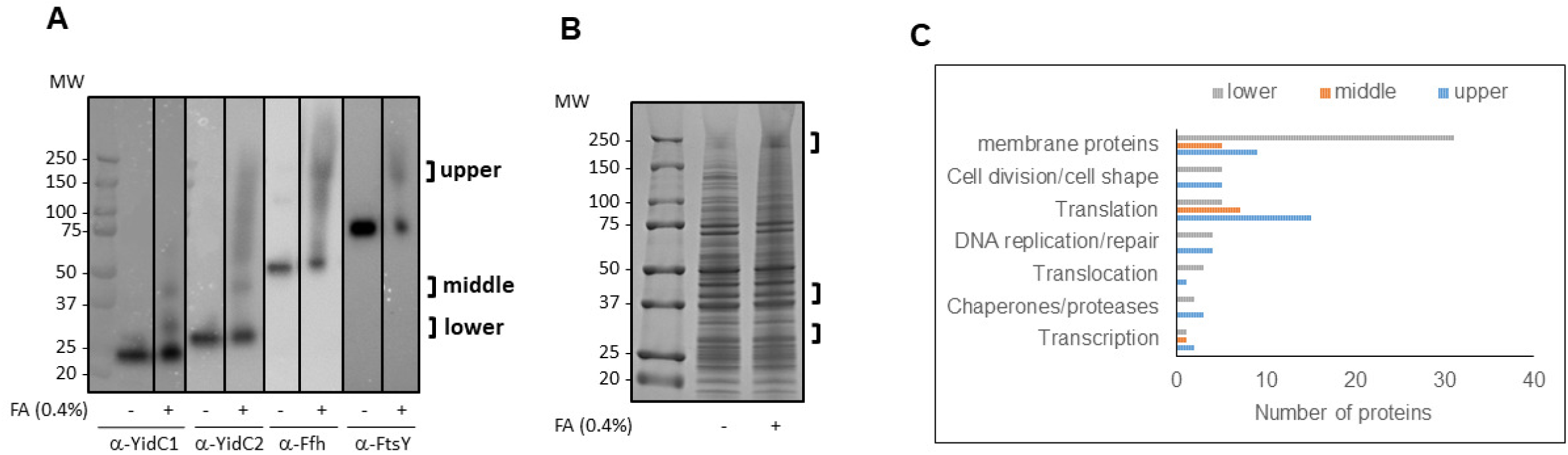
Formaldehyde cross-linking of *S. mutans* results in gel shifts of protein translocation machinery components present in whole cell lysates. **(A)** Whole cell lysates of untreated (-) *S. mutans* strain UA159 or cells treated with 0.4% formaldehyde (+) were analyzed by Western blot using anti-YidC1, anti-YidC2, anti-Ffh and anti-FtsY antibodies. Brackets indicate regions of reactivity subjected to further analysis. Bands corresponding to YidC1, (24 kDa), YidC2 (27 kDa), Ffh (54 kDa), and Ftsy (75 kDa) are apparent in untreated and formaldehyde cross-linked samples. **(B**) Corresponding Coomassie blue stained SDS-polyacrylamide gel indicating location of excised gel slices sent for mass spectrometry analysis. **(C)** Histogram showing number of proteins in indicated categories in upper, middle and lower excised gel slices.

A total of 65, 38, and 119 proteins were identified in the upper, middle, and lower molecular weight gel slices, respectively, of the cross-linked but not non-cross-linked sample (Table S1). The lower region, recognized by anti-YidC1 antibodies, contained the highest proportion of membrane proteins (31/119). These may therefore represent substrates of a pathway that involves YidC1, but not YidC2. In contrast, the higher region recognized by anti-Ffh, anti-FtsY, and anti YidC2 antibodies, and the middle region recognized by anti-YidC1 and anti-YidC2 antibodies, contained fewer membrane proteins, 9/65 and 5/38, respectively. Most of the other non-integral membrane proteins identified in all three regions had previously been found to be membrane-associated in our proteomic analysis of membrane preparations derived from protoplasts of *S. mutans* wild-type compared to *ΔyidC1, ΔyidC2, Δffh*, and *Δffh/yidC1* mutants [10]. Many membrane-associated proteins are components of multimeric membrane-localized complexes that also contain integral membrane components. Thus, identification of membrane-associated proteins may indirectly reflect the actual integral membrane substrates. We also identified multiple proteins involved in DNA replication and repair, transcription, translation, and cell division suggesting an extensive coordinated cellular machinery that includes membrane protein translocation (Table 1).

**Table 1:** Summary of proteins present in upper, middle, and lower molecular weight regions, identified by Western blot gel shift assays of whole cell lysates of formaldehyde cross-linked *S. mutans* detected with anti-YidC1, YidC2, Ffh, and FtsY-specific antibodies

| **Upper Region:** reactive with anti-YidC2, Ffh and FtsY antibodies | **Middle Region:** reactive with YidC1 and YidC2 | **Lower Region:** reactive with anti-YidC1 antibodies |
| --- | --- | --- |
| **Translation** |  |  |
| 50S ribosomal protein L1 (SMU_1626)* | 50S ribosomal protein L1 (SMU_1626)* | Putative ribosomal protein S1; sequence specific DNA-binding protein (SMU_1200)<sup>§</sup> |
| 50S Ribosomal Protein L2 (SMU_2167)* | 50S Ribosomal Protein L2 (SMU_2167)* | Dimethyladenosine transferase, 16S rRNA methyltransferase (SMU_349) |
| 50S ribosomal protein L7/L12 (SMU_960) | 50S ribosomal protein L5 (SMU_2015) | Putative tRNA pseudouridine synthase A (SMU_84) |
| Putative ribosomal protein S1; sequence specific DNA-binding protein (SMU_1200)<sup>§</sup> | 50S ribosomal protein L18 (SMU_2010) | Translation elongation factor EF-Tu (SMU_714)<sup>§</sup> |
| 30S ribosomal protein S3 (SMU_2021)* | 30S ribosomal protein S2 (SMU_2032) | Conserved hypothetical protein, 16S rRNA methyltransferase (SMU_1659c) |
| 30S ribosomal protein S5 (SMU_2009) | 30S ribosomal protein S3 (SMU_2021)* |  |
| 30S ribosomal protein S10 (SMU_2026c) | 30S ribosomal protein S13 (SMU_2003) |  |
| 30S ribosomal protein S11 (SMU_2002) |  |  |
| 30S ribosomal protein S18 (SMU_1858) |  |  |
| Translation elongation factor EF-Tu (SMU_714)<sup>§</sup> |  |  |
| Translation elongation factor G (SMU_359) |  |  |
| Putative translation elongation factor TS (SMU_2031) |  |  |
| Translation initiation factor 2 (SMU_421) |  |  |
| Putative translation initiation factor IF3 (SMU_697) |  |  |
| Putative alanyl-tRNA synthetase (alanine--tRNA ligase) (SMU_650) |  |  |
| **Translocation** |  |  |
| Preprotein translocase subunit SecA (SMU_1838) |  | Putative preprotein translocase SecY protein (SMU_2006) |
|  |  | Putative secreted protein, preprotein translocase subunit YajC (SMU_1787c) |
|  |  | Putative signal peptidase II (SMU_1661c) |
| **Chaperones/proteases** |  |  |
| Peptidyl-prolyl isomerase RopA (trigger factor) (SMU_91)<sup>§</sup> |  | Peptidyl-prolyl isomerase RopA (trigger factor) (SMU_91)<sup>§</sup> |
| Heat shock protein. DnaK (HSP-70) (SMU_82) |  | Putative chaperonin GroEL (SMU_1954)<sup>§</sup> |
| Putative chaperonin GroEL (SMU_1954)<sup>§</sup> |  |  |
| **Cell division/cell shape** |  |  |
| Putative septation ring formation regulator (SMU_1276c) |  | Putative cell shape-determining protein MreC (SMU_20) |
| Putative cell division protein DivIVA (SMU_557) |  | Putative cell division protein RodA (SMU_1279c) |
| Putative cell division protein FtsZ (SMU_552) |  | Integral membrane protein possibly involved in D-alanine export, D-alanyl-lipoteichoic acid biosynthesis protein DltB (SMU_1690) |
| Cell division protein FtsA (SMU_551) |  | Putative cell division protein FtsH (SMU_15)<sup>§</sup> |
| Putative cell division protein FtsH (SMU_15)<sup>§</sup> |  | Conserved hypothetical protein cell division protein FtsW (SMU_172) |
| **DNA replication/repair** |  |  |
| DNA-directed RNA polymerase, alpha subunit (SMU_2001) |  | Putative type II restriction endonuclease (SMU_506) |
| DNA-dependent RNA polymerase, beta subunit (SMU_1990) |  | Putative site-specific DNA-methyltransferase (SMU_504) |
| DNA-dependent RNA polymerase, beta' subunit (SMU_1989) |  | Putative endonuclease III (DNA repair) (SMU_1650) |
| Recombination protein RecA (SMU_2085) |  | Putative DNA polymerase III, delta subunit (SMU_1662) |
| **Transcription** |  |  |
| DNA-dependent RNA polymerase sigma subunit; major sigma factor (sigma 70/42) (SMU_822) | DNA-dependent RNA polymerase. beta' subunit (SMU_1989) | Conserved hypothetical protein, DNA-directed RNA polymerase subunit delta (SMU_1936c) |
| Putative tRNA isopentenylpyrophosphate transferase, tRNA dimethylallyltransferase (SMU_1477) |  |  |
| **Putative substrates (proteins with one or more predicted transmembrane domains)** |  |  |
| Putative septation ring formation regulator (SMU_1276c) | Hypothetical protein (SMU_591c)* | Conserved hypothetical protein peptide ABC transporter substrate-binding protein (SMU_1447c) |
| Putative ABC transporter, permease protein (SMU_1007) | Serine protease HtrA (SMU_2164) | Putative carbonic anhydrase precursor (SMU_1595) |
| Putative PTS system, fructose-specific enzyme IIABC component (SMU_872) | Putative PTS system, mannose-specific component IID (SMU_1879) | Putative deacetylase (SMU_623c) |
| Hypothetical protein (SMU_591c)* | Hemolysin (SMU_1693) | Conserved hypothetical protein, glycosyl transferase (SMU_834) |
| Putative ABC transporter, ATP-binding protein ComA (SMU_286) | Hypothetical protein APQ13_07375 (ParE_toxin) (SMU_40) | Putative PTS system, glucose-specific IIABC component (SMU_2047)<sup>§</sup> |
| Conserved hypothetical protein, protease (SMU_235) |  | Hypothetical protein (SMU_832) |
| Putative cell division protein FtsH (SMU_15)<sup>§</sup> |  | Putative amino acid ABC transporter, periplasmic amino acid-binding protein (SMU_933) |
| Putative PTS system, glucose-specific IIABC component (SMU_2047)<sup>§</sup> |  | Conserved hypothetical protein, cyclic nucleotide-binding protein (SMU_1307c) |
| Putative PTS system, trehalose-specific IIABC component (SMU_2038) |  | Putative cell shape-determining protein MreC (SMU_20) |
|  |  | Putative glycosyltransferase (SMU_833) |
|  |  | Conserved hypothetical protein (SMU_1477c) |
|  |  | Putative undecaprenyl-phosphate-UDP-MurNAc-Pentapeptide transferase (SMU_456) |
|  |  | Putative cell division protein RodA (SMU_1279c) |
|  |  | Integral membrane protein possibly involved in D-alanine export, D-alanyl-lipoteichoic acid biosynthesis protein DltB (SMU_1690) |
|  |  | Putative transmembrane protein, permease OppC (SMU_257) |
|  |  | Conserved hypothetical protein (SMU_1111c) |
|  |  | Putative amino acid permease (SMU_1450) |
|  |  | Hypothetical protein (SMU_503c) |
|  |  | Hypothetical protein (SMU_1249c) |
|  |  | Putative preprotein translocase SecY protein (SMU_2006) |
|  |  | Putative sodium/amino acid (alanine) symporter (SMU_1175) |
|  |  | Putative glycosyl transferase N-acetylglucosaminyltransferase), RgpG (SMU_246) |
|  |  | Putative secreted protein, preprotein translocase subunit YajC (SMU_1787c) |
|  |  | Putative endolysin, N-acetylmuramoyl-L-alanine amidase (SMU_707c) |
|  |  | Putative manganese transporter (SMU_770c) |
|  |  | Putative serine/threonine protein kinase (SMU_484) |
|  |  | Putative cell division protein<br>FtsH (SMU_15)<sup>§</sup> |
|  |  | Hypothetical protein<br>(SMU_1161c) |
|  |  | Putative drug-export protein;<br>multidrug resistance protein,<br>XRE family transcriptional<br>regulator (SMU_745) |
|  |  | Putative ABC transporter,<br>periplasmic ferrichrome-<br>binding protein (SMU_998) |
|  |  | Cell wall-associated protein<br>precursor WapA (SMU_2159) |

\*indicates proteins present in both upper and middle molecular weight gel slices.
<sup>§</sup>indicates proteins detected in regions reactive with both anti-YidC1 and anti-YidC2 antibodies.

That the upper gel-shifted region was recognized by anti-FtsY, anti-Ffh and anti-YidC2, but not anti-YidC1, antibodies, suggests that YidC2, but not YidC1, likely works in concert with the SRP pathway. The middle and lower regions were recognized by both anti-YidC1 and anti-YidC2 antibodies, or only by anti-YidC1 antibody, respectively, but not by anti-Ffh or anti-FtsY antibodies. This result suggests that the SRP pathway does not normally interact with YidC1. Because *S. mutans* survives deletion of *yidC2*, but not of *yidC1* and *yidC2* [7], YidC1 may cooperate with the SRP pathway only when YidC2 is absent. Our previous membrane proteomic analysis of *S. mutans* protein transport mutants suggested that the SRP pathway acts in concert with at least one YidC paralog in the insertion of multiple substrates [10]. However, that study utilized deletion mutants while the current study is more indicative of protein interactions in the wild-type strain. The identification of SecY and YajC, as well as YidC1, in the lower region suggests that these represent components of a pathway that operates independently of YidC2 and/or the SRP, and supports previous reports of a SecY-YidC interaction in *E. coli* that is modulated by YajC [15, 17, 36, 41]. Thus in *S. mutans*, YidC1 likely serves as the major interaction partner of SecY/YajC. It has also been reported in *E. coli* that SecYEG and YidC compete for binding to the SRP receptor, FtsY [17]. Our results suggest, therefore, that YidC1 and YidC2 participate in two distinct pathways such that YidC1 functions in concert with the SecY translocon independently of the SRP pathway, while YidC2 functions primarily in concert with the SRP pathway. It is interesting that the membrane-associated SecA molecular motor of the general secretion pathway [42, 43] was identified in the upper region recognized by anti-Ffh, anti-FtsY, and anti YidC2 antibodies, but not in the lower region that contained SecY and was recognized by anti-YidC1 antibodies. One reason for detection of SecA in a section of the gel reactive with anti-SRP antibodies may be through indirect bridging via ribosomal proteins. SecA and the SRP have been shown to bind to the same location on the *E. coli* ribosome in order to sort cellular proteins into distinct pathways for secretion through or insertion into the membrane at the site of translation [44]. Our experiment utilized whole cell lysates and was therefore biased towards identification of cytoplasmic or membrane proteins and not expected to identify secreted SecA substrates, although SecA itself would be in proximity of SRP components on the ribosome. Also of interest, the middle region reactive with both anti-YidC1 and anti-YidC2 antibodies, did not include SecY, YajC, or SRP components. *E. coli* YidC is known to function in a YidC only pathway in the insertion of small membrane proteins [13, 22, 45]. Interestingly, 4 out of the 5 membrane proteins with only 1 or 2 TM domains were present in the middle region (Table 1). This suggests that YidC1 and/or YidC2 can work independently of SecY, and the SRP pathway, in the insertion of a limited number of small membrane proteins.

In an attempt to identify respective substrates of YidC1-SecY, YidC1/2, or SRP-YidC2 mediated pathways, those proteins predicted to have one or more transmembrane were categorized according to their presence in upper, middle, or lower regions of the gel (Table 1). The identification of a higher proportion of membrane proteins in the lower region suggests that YidC1-SecY/YajC is a widely used pathway for membrane protein insertion. That is, YidC1 likely represents the “housekeeping” paralog in *S. mutans.* A closer examination of membrane proteins in the lower region revealed known or putative metal transporters including SMU_770C, SMU_998, and a putative zinc ABC transporter ATP-binding protein (SMU_1994) suggesting that these particular metal transporters utilize the YidC1-SecY/YajC pathway for insertion. Also, proteins encoded in an operon of unknown function that includes SMU_832, SMU_833, and SMU_834 were identified in the lower region. Proteins in the upper region, suggestive of insertion by a coordinated SRP-YidC2 pathway, included multiple sugar transporters and several ABC transporters including the competence-associated protein ComA. *S. mutans* deletion mutants lacking *ffh* or *yidC2* exhibit impaired genetic competence, but this property is less impacted by elimination of *yidC1* [46]. We did not observe a preference for single TM compared to multi-pass membrane proteins for the YidC1-SecY/YajC or the SRP-YidC2 pathways. In contrast, as stated above, membrane proteins from the middle region that may represent substrates of a YidC1 and/or YidC2 only pathway were mostly single or double-pass membrane proteins except for hemolysin (SMU_1693), which contains 4 predicted TM domains.

While *in vivo* whole cell cross-linking identified a relatively low number of potential membrane-localized substrates of the putative SRP-YidC2 mediated protein translocation pathway, a high proportion of proteins (30/65) present in the upper region of the gel are involved in DNA replication/repair, transcription, translation, and cell division/cell shape (Table 1). These results support the idea that the SRP-YidC2 co-translational translocation pathway operates in the context of a larger consortium of proteins that make up an integrated higher order machinery, which couples replication, transcription, and cell division with membrane insertion of a subset of membrane proteins. In contrast, approximately 25% of proteins present in the lower gel slice representative of the putative SecY-YajC /YidC1 pathway are membrane proteins. Additionally, multiple proteins identified in this region are associated with replication, transcription, or translocation. The streptococcal homolog of the ribosome-associated chaperone trigger factor, RopA, was identified in both the upper and lower regions that likely represent the SRP-YidC2 and SecY-YidC1-mediated protein translocation pathways, respectively. Trigger factor has been reported to bind to the same ribosomal protein at the peptide exit site as the SRP pathway [47, 48]; therefore, the finding of RopA and Ffh in the same gel slice was not unexpected.

Because a whole cell cross-linking approach is limited by the accessibility of exposed functional groups in the target proteins to formaldehyde [49], the actual integral membrane substrates of the insertion machinery were likely underrepresented in our dataset due largely to their being buried within the membrane and inaccessible to the cross-linking reagent. Surprisingly YidC1, YidC2, Ffh, and FtsY themselves were also not identified in either cross-linked or non-cross-linked samples, although they were clearly present as evidenced by their detection by Western blot and migration at the correct molecular weight. It is possible that they were not amenable to or present in sufficient quantity for detection by mass-spectrometry. Western blot with high quality antibodies can be more sensitive than standard bottom-up MS in the detection of certain proteins of low abundance [50]. To overcome this potential limitation, we also employed immunocapture experiments in an attempt to improve sensitivity.

### Dynabead™ immunocapture of protein complexes from *S. mutans* lysates using anti-YidC2 antibodies

To identify potential binding partners of YidC2 and to characterize this insertase’s interactome, an immunocapture approach was undertaken in which anti-YidC2 antibodies were covalently coupled to magnetic Dynabeads™. The polyclonal rabbit antibodies used were made against synthetic peptides corresponding to the YidC2 C-terminal tail and cytoplasmic loop between TM2 and TM3. Whole cell lysates from untreated and formaldehyde-treated cells of *S. mutans* strain NG8, and its corresponding *ΔyidC2* mutant, were reacted with the antibody-coupled beads and bound proteins were eluted with glycine-HCl, pH 2.0. Aliquots of each sample were analyzed by Western blot (Fig. 2A). As expected, a 27 kDa YidC2 band was identified in the wild-type (WT), but not *ΔyidC2* strain (Fig. 2B). An additional band reactive with anti-YidC2 antibodies was also observed in the cross-linked sample from the WT, but not samples from the mutant strain, or the non-cross-linked control sample from the WT strain. Other higher molecular weight bands were observed in a replicate negative control blot probed only with goat-anti-rabbit heavy chain specific secondary antibodies. These represent anti-YidC2 antibodies present in the eluate that leached from the coupled Dynabeads™ (Fig. 2C). Gel slices corresponding to the ∼45 kDa region of interest identified by Western blot were cut from SDS-PAGE gels of all four samples and analyzed by MS. A total of 269 proteins were identified in the WT cross-linked sample, 229 in the WT non-cross-linked sample, 246 proteins in the *ΔyidC2* cross-linked sample, and 284 proteins in the *ΔyidC2* non-cross-linked sample. Sixty-eight proteins were present in the WT cross-linked sample and 31 in the non-cross-linked sample that were absent from the corresponding *ΔyidC2* samples (summarized in Table S2). Six of those were shared between cross-linked and non-cross-linked samples. The presence of such shared proteins may represent direct binding partners of YidC2 that do not need to be cross-linked to be co-captured with it. A graphical representation of the types of proteins co-captured with YidC2 is shown (Fig. 2D).

**Fig. 2.**
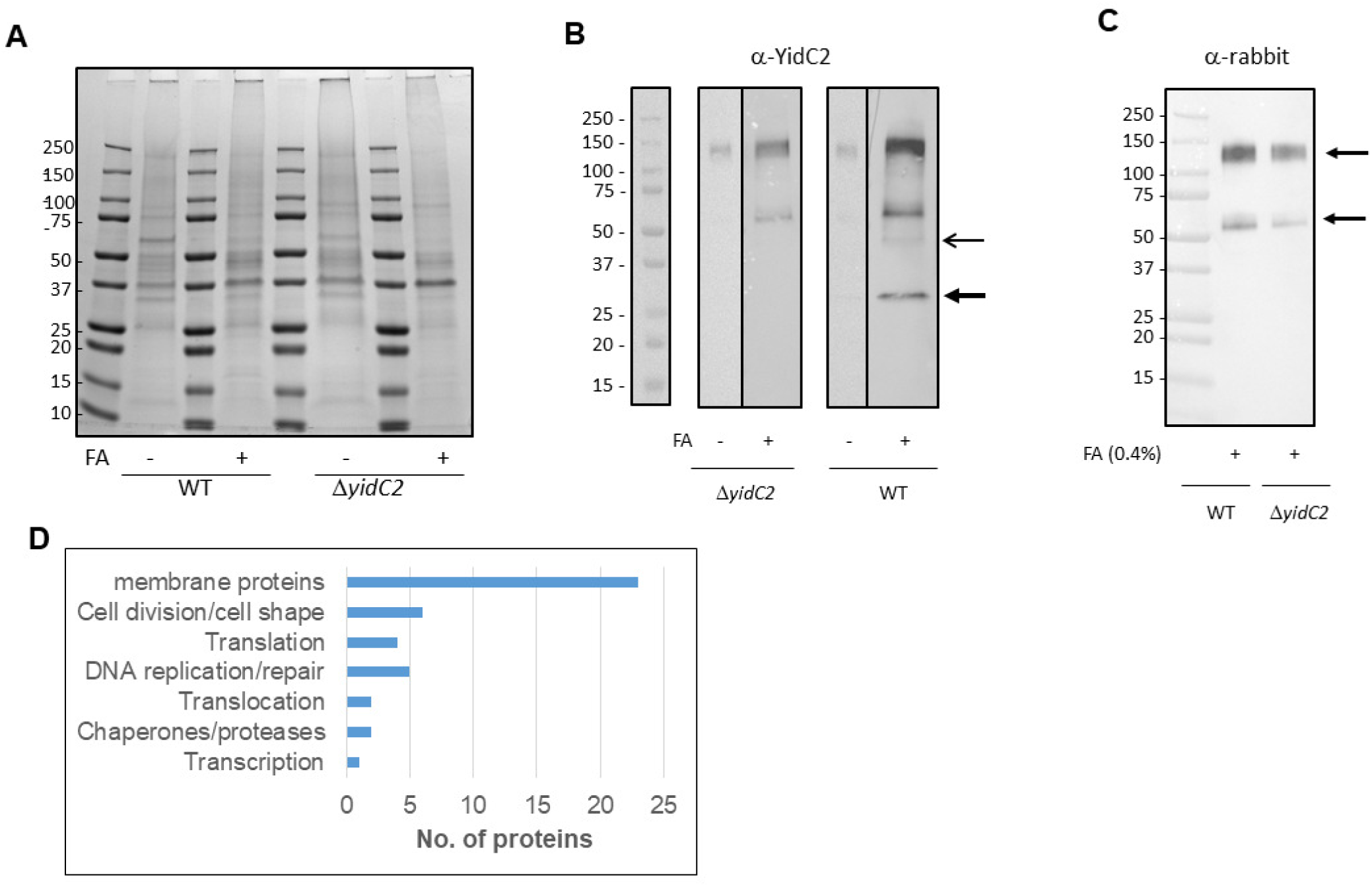
Immunocapture of YidC2 and associated protein complexes from whole cell lysates (WCL) of *S. mutans* using anti-YidC2 antibodies coupled to Dynabeads™. **(A)** SDS-PAGE. Dynabeads™ conjugated with anti-YidC2 antibodies were reacted with whole cell lystaes from untreated (-) or 0.4% formaldehyde cross-linked (+) wild-type *S. mutans* strain NG8 (WT) or corresponding *ΔyidC2* mutant and eluted with 0.5 N NH_4_OH, 0.5 mM EDTA. Migration of molecular weight standards is indicated. **(B)** Western blot of samples shown in (A). Thick arrow indicates YidC2. Thin arrow indicates the gel-shifted band seen only in the cross-linked sample from the WT strain. This region was excised for each of the four samples from the Coomassie blue gel, stained, and subjected to mass spectrometry analysis. **(C)** A replicate negative control Western blot probed with goat anti-rabbit secondary antibodies identifies the migration of anti-YidC2 antibodies that leached from the column during the elution step. **(D)** Histogram showing the types of proteins co-captured with YidC2 from the WT strain.

Twenty-four of the 99 total immunocaptured proteins identified in WT but not *ΔyidC2* samples are predicted to have one or more TM domains, with the rest having been identified as membrane-associated in our previous membrane proteomic analyses [10] (Table S2). Potential integral membrane substrates of YidC2 identified by this immunocapture experiment include two subunits of the PTS mannose transporter, metalloprotease (RseP), histidine kinases, enzymes, and cell wall/cell division related proteins (Table 2). In agreement with our prior gel shift experiment and analysis of the upper gel slice, we identified SecA as well as other proteins involved in DNA replication/repair, transcription, translation, and cell division/cell shape in associated with YidC2 (Fig. 2D, Panel D). Again this suggests that translocation is part of a coordinated machinery that incorporates additional processes beyond protein translation. In contrast to the gel shift assay, in which both YajC and SecY were detected in the lower gel slice reactive with anti-YidC1 antibodies, in the current immunocapture experiment YajC, but not SecY, was identified as part of YidC2 interactome. Certain differences in the apparent YidC2 interactome identified following Western blot gel shift, as opposed to Dynabead™ immunocapture, may relate to those domains of YidC2 available for binding to substrates or other proteins during the two experimental approaches. Antibodies against cytoplasmic loop 1 and the YidC2 tail domain were utilized for Dynabead™ capture of YidC2 and associated proteins. Thus binding of proteins that interact specifically with either of these regions may have prevented efficient capture of YidC2 by the antibody-coupled magnetic beads. That is, the immunocapture dataset was likely biased against proteins that react with cytoplasmic loop 1 or the YidC2 C-terminal tail. In *E. coli*, cytoplasmic loop 1 of YidC has been reported to interact with SecY in an *in vivo* photo-crosslinking assay [17]. Thus, occupancy of the corresponding loop in YidC2 by SecY could potentially have blocked reactivity with the anti-YidC2 loop antibody and explain why YajC, but not SecY, was identified in the immunocapture assay. Unlike the Western blot gel shift results, we did not identify Ffh or FtsY in association with YidC2 in the immunocapture experiment. If the cooperative activity of YidC2 with the SRP pathway depends on an interaction mediated by its C-terminal tail, that could preclude its efficient capture by anti-tail-specific antibodies. Previous domain swapping experiments support this conjecture in that stress tolerance was complemented in a Δ*yidC2* background with chimeric YidC1 whose C-terminal tail was replaced with that of YidC2 [8]. Collectively, our anti-YidC2 immunocapture assay identified not only the translocation machinery components SecA and YajC, but also ribosomal proteins, chaperones and proteases, enzymes involved in DNA replication and repair, and proteins responsible for cell wall generation and cell division. Because of issues with low coupling efficiency of the anti-YidC2 antibodies to Dynabeads™ and the problem with antibody leaching, this approach was not attempted with anti-YidC1 antibodies.

**Table 2:**
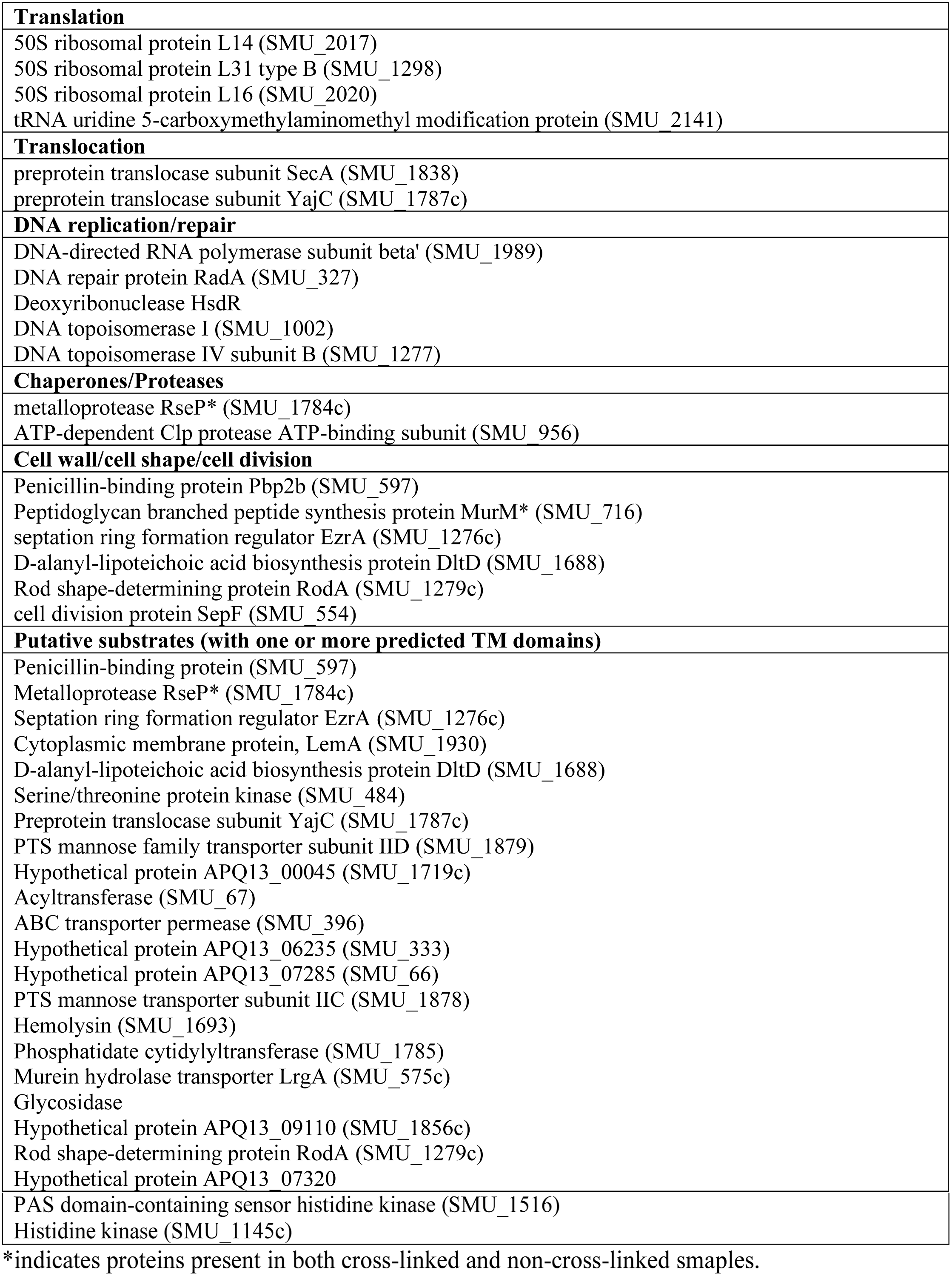
List of proteins belonging to various functional categories uniquely present in the formaldehyde cross-linked and non-cross-linked *S. mutans* wild-type (WT) whole cell lysates, but absent from the *ΔyidC2* mutant strain by immunocapture with anti-YidC2 antibodies.

### Difference gel electrophoresis of *S. mutans* proteins captured by YidC1 or YidC2 C-terminal tails

As an alternative to immobilization of anti-YidC antibodies to Dynabeads™, we also utilized a Glutathione-S-transferase (GST)-tagged based pull-down approach. While it is difficult to express *S. mutans yidC1* and *yidC2* in *E. coli* to sufficient levels for large scale protein purification, both the YidC1 and YidC2 C-terminal tails are soluble, and easily tagged and purified. We constructed fusion proteins of the YidC1 and YidC2 C-terminal domains with GST and affinity purified the recombinant polypeptides on Glutathione Sepharose™ (Fig. S1). Because domain swapping experiments have demonstrated that the positively-charged tails of *S. mutans* YidC1 and YidC2 contribute to certain functional attributes of each paralog [8], we expected a subset of YidC1 and YidC2 binding partners to interact with these domains. After purification, the GST-tagged YidC1/2-tail fusion proteins were reacted with *S. mutans* whole cell lysates (non-cross-linked) and captured on immobilized glutathione using GST as a negative control. Following elution with reduced glutathione the three samples were individually labeled with a different CyDye fluorescent dye and subjected to 2D-difference gel electrophoresis (DIGE) (Fig. 3). One hundred and twenty-one spots were identified as being captured by GST-YidC1CT and/or GST-YidC2CT, but not by GST (Fig. S2). A complete list of all proteins identified in each of the gel spots is shown in Table S3. A summary of the proteins pulled down with GST-Yid1CT (green spots), GST-YidC2CT (red spots), or both (yellow spots) is shown in Table S4. Seventy-four proteins were co-captured with GST-YidC1CT, and 37 with GST-YidC2CT (Table S3). Of those, 42 were uniquely co-captured with GST-YidC1CT, while only 5 were uniquely co-captured with GST-YidC2CT (Table S4).

**Fig. 3.**
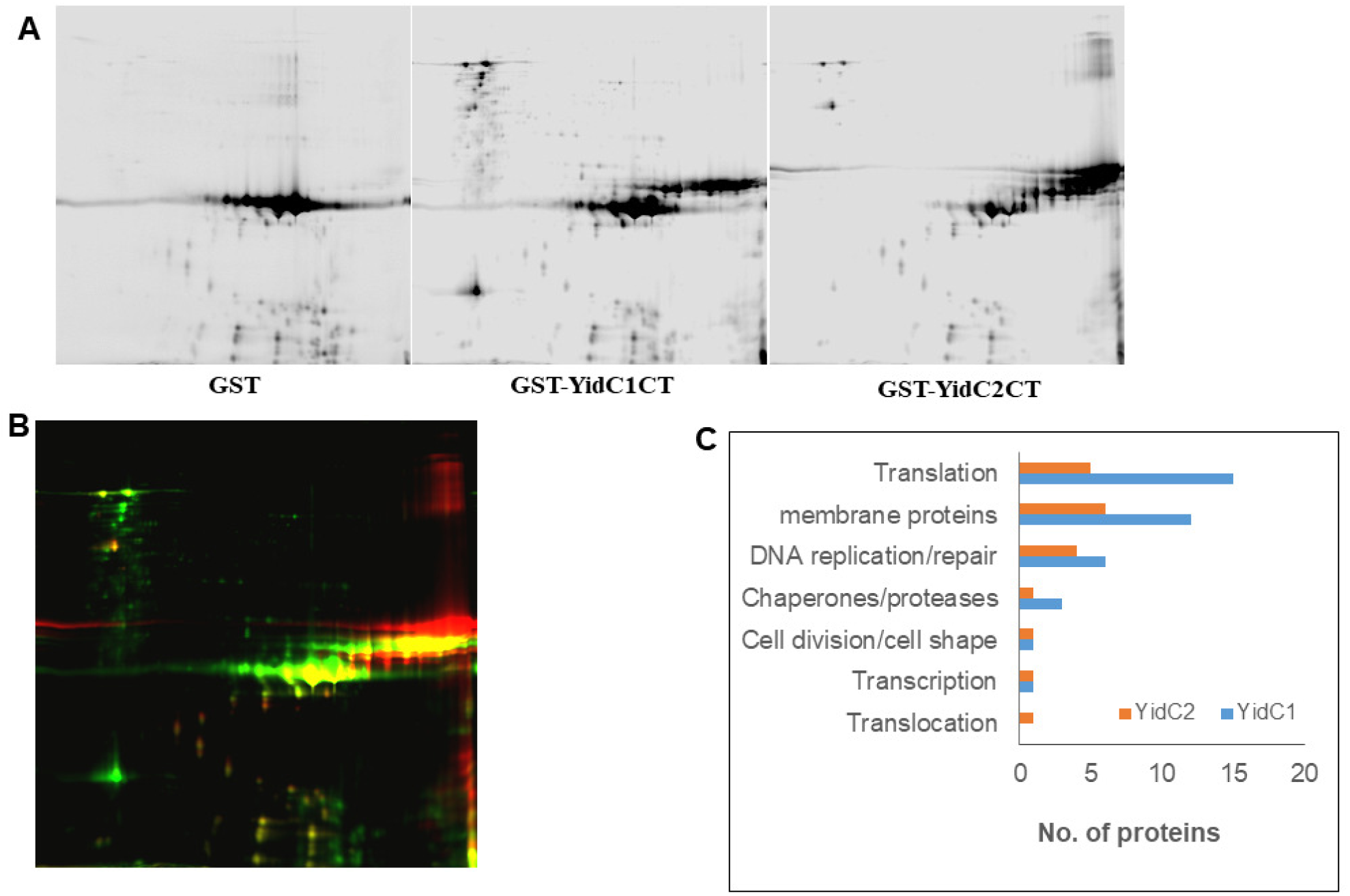
Proteins co-captured with GST, GST-YidC1CT or GST-YidC2CT analyzed by 2D-DIGE. **(A)** *S. mutans* whole cell lysates were reacted with the indicated GST polypeptide and captured using glutathione affinity chromatography. The eluted samples were labeled with CyDye DIGE fluors (YidC1CT with red Cy3, YidC2CT with green Cy2, and GST with blue Cy5), and separated on a single 2D gel, by isoelectric focusing in the first dimension and SDS-PAGE in the second dimension. Black and white images for each sample are shown. **(B)** Signals from each dye were scanned and the three images overlaid. One hundred and twenty separate spots (shown in Fig. S2) were excised from the gel for mass spectrometry analysis. **(C)** The numbers and types of proteins associated with GST-YIDC1CT compared to GST-YidC2CT are shown.

The types of proteins co-captured with GST-YidC1CT compared to GST-YidC2CT are summarized in Table 3. Proteins with 1 or more transmembrane domains were considered as putative substrates. Eleven different integral membrane proteins were found as part of the YidC1-tail interactome, including 5 that were also pulled down with GST-YidC2CT. Most of these were transporters with the exception of the cell division protein FtsH, and a histidine kinase (SMU_486). All non-integral membrane proteins identified by DIGE (∼85%) had previously been identified as membrane-associated during proteomic analysis of *S. mutans* protoplast-derived membrane preparations [10]. The predominance of non-integral membrane proteins in the DIGE dataset suggest that the YidC1 and YidC2 C-terminal tails do not play a prominent role in recognizing and binding substrates. Twenty of the 24 membrane-associated proteins from the YidC2-tail interactome were also co-captured with GST-YidC1CT. Interestingly, the SRP component protein Ffh was found in association with the YidC2-tail, but not with the YidC1-tail. This supports data from the *in vivo* cross-linking experiments that suggested a cooperative SRP-YidC2 pathway, and explains why appending the YidC2 tail onto YidC1 enables the chimeric protein to ameliorate the *ΔyidC2* phenotype. Presumably this manipulation allows the YidC1 insertase to interact with Ffh and function in concert with the SRP pathway machinery. None of the components of the SecYEG translocon, nor YajC, were identified in association with either of the C-terminal tails, thus these domains likely do not contribute to YidC1 or YidC2 interactions with the translocon itself.

**Table 3:** List of *S. mutans* proteins pulled down with GST-YidC1CT and/or GST-YidC2CT, but not GST, identified by 2D-DIGE and mass spectrometry

| **GST-YidC1-CT** | **GST-YidC2-CT** |
| --- | --- |
| **Translocation** |  |
|  | Signal recognition particle protein (SMU_1060) |
| **Translation** |  |
| 50S Ribosomal Protein L2 (SMU_2160)<br>50S ribosomal protein L6 (SMU_2011)<br>50S ribosomal protein L13 (SMU_169)<br>Putative ribosomal protein S1 (SMU_1200)<br>30S ribosomal protein S2 (SMU_2032)<br>30S ribosomal protein S4 (SMU_2135c)<br>30S ribosomal protein S7 (SMU_358)<br>30S ribosomal protein S8 (SMU_2012)<br>30S ribosomal protein S17 (SMU_2017)<br>30S ribosomal protein S21 (SMU_818)<br>Elongation factor Tu (SMU_714)<br>Phenylalanyl-tRNA synthetase subunit alpha (SMU_1512)<br>Glycyl-tRNA synthetase subunit alpha (SMU_445)<br>S1 RNA-binding domain-containing protein (smu_1623c)<br>tRNA (adenine(22)-N(1))-methyltransferase (SMU_1464c) | 50S Ribosomal Protein L2 (SMU_2160)<br>50S ribosomal protein L6 (SMU_2011)<br>50S ribosomal protein L13 (SMU_169)<br>30S ribosomal protein S7 (SMU_358)<br>30S ribosomal protein S17 (SMU_2017) |
| **DNA replication/repair** |  |
| DNA polymerase III, gamma/tau subunit (SMU_1581)<br>DNA polymerase III PolC (SMU_123)<br>DNA-directed RNA polymerase subunit alpha (SMU_2001)<br>DNA polymerase I (POL I) (SMU_297)<br>DNA repair protein RecN (SMU_585)<br>DNA mismatch repair protein MutS (SMU_2091c) | DNA polymerase III, gamma/tau subunit (SMU_1581)<br>DNA polymerase III PolC (SMU_123)<br>DNA-directed RNA polymerase subunit alpha (SMU_2001)<br>Holliday junction-specific endonuclease (SMU_469) |
| **Transcription** |  |
| Probable DNA-directed RNA polymerase subunit delta (SMU_96) | Probable DNA-directed RNA polymerase subunit delta (SMU_96) |
| **Chaperones/Proteases** |  |
| molecular chaperone DnaK (SMU_82)<br>Chaperone protein ClpB (SMU_1425)<br>heat shock protein GrpE (SMU_81) | Molecular chaperone DnaK (SMU_82) |
| **Cell wall/cell shape/cell division** |  |
| cell division protein FtsH (SMU_15) | cell division protein FtsH (SMU_15) |
| **Putative substrates (with one or more predicted TM domains)** |  |
| Cell division protein FtsH (SMU_15)<br>Putative amino acid ABC transporter, permease protein (SMU_1216c)<br>ABC transporter ATP-binding protein (SMU_906)<br>Histidine kinase (SMU_486)<br>Putative PTS system, glucose-specific IIABC (SMU_2047)<br>Putative ABC transporter, substrate-binding protein (SMU_651c)<br>Dextranase (SMU_2042)<br>Hypothetical protein (SMU_791c)<br>Conserved hypothetical protein (SMU_485)<br>Potassium transporter peripheral membrane protein (SMU_1708)<br>EamA family transporter (SMU_1560) | ABC transporter ATP-binding protein (SMU_906)<br>Putative ABC transporter, substrate-binding protein (SMU_651c)<br>Histidine kinase (SMU_486)<br>Cell division protein FtsH (SMU_15)<br>Putative amino acid ABC transporter, permease protein (SMU_1216c) |

As described above in the gel shift and YidC2 immunocapture experiments, numerous ribosomal proteins, as well as other components of the translation machinery, were captured in association with YidC1 and/or YidC2. Such proteins were found irrespective of whether Ffh and FtsY were also present, suggesting that either *S. mutans* YidC paralog can act to support co-translational protein translocation in the absence of the SRP pathway. Indeed, YidC2 was previously demonstrated to complement Oxa1 deficiency in yeast mitochondria that lack an SRP pathway [27]. While YidC1 was present in yeast cell extracts, this paralog was not properly imported into the mitochondria and therefore could not be assessed in complementation experiments[27]. When overexpressed in *E.coli*, both YidC1 and YidC2 of *S. mutans* were found to interact with translating and non-translating ribosomes by a tail-dependent mechanism [51]. In the current study, the large ribosomal subunit protein, L2, was the most abundant ribosomal protein pulled down by both the GST-YidC1CT and GST-YidC2CT fusion polypeptides. In *E. coli*, L2 not only acts as a structural component of the ribosome, it is also processed to a truncated derivative (tL2) that can interact with the RNA polymerase alpha subunit and modulate transcription [52]. *E. coli* L2 has also been reported to interact with the Hsp90 homolog HtpG to modulate its ATPase activity, and also to bind to other chaperones including DnaK/DnaJ/GrpE and GroEL/GroES [53]. Full-length and truncated *E. coli* L2 also interact with DnaA to modulate DNA replication [54]. DNA encoding *S. mutans* L2 and rL2 was cloned and the recombinant his-tagged proteins were tested by ELISA to determine whether either form interacts directly with the C-terminal tails of YidC1 or YidC2. Neither L2 nor rL2 demonstrated significant binding to GST-YidC1CT or to GST-YidC2CT (Fig. S3). Likewise, SecA, which had been observed in conjunction with YidC2 in both Western blot gel-shift and immunocapture experiments, did not react directly with GST-YidC2CT (or GST-YidC1CT) (Fig. S3). This suggests that the association of SecA with YidC2 is indirect, or nor mediated by the YidC2 tail.

Similar to the previous experiments, GST-YidC1CT and GST-YidC2CT also captured a variety of proteins including chaperones and those involved in replication, transcription, translation, and cell division/cell shape again suggesting that all these processes are temporally and spatially connected. These data are consistent with the identification of coupled transcription/translation in other bacteria, which may also integrate aspects of DNA replication [55-59]. Both YidC1 and YidC2 contribute to proper cell wall biosynthesis and cell morphology in *S. mutans* [9], thus capture of proteins in this category is consistent with previously described mutant phenotypes.

### Determination of YidC1 and YidC2 interactomes and functional annotation

When proteins from all experiments were evaluated in composite, 88 were identified as being associated with both YidC1 and YidC2, while 123 or 131 were uniquely associated with YidC1 or YidC2, respectively (Fig. 4A). When possible, proteins were assigned to functional categories by DAVID analysis (Fig. 4B). The most prevalent functional category in both interactomes was transferase. Functional annotation also shows that the YidC2, compared to YidC1, interactome was enriched in a number of functional categories including ATP-binding proteins, metalloproteins, carbon metabolism, oxidoreductases, cell division, GTPase activity, and branched chain amino acid pathways. This may explain why the phenotypic consequence of elimination of YidC2 is far more pronounced than elimination of YidC1 [7, 8, 46]. In contrast, the only instances in which the YidC1 interactome equaled or exceeded that of YidC2 were in the transferase, and purine and pyrimidine metabolism categories. Of note, however, a greater number of proteins in the YidC1 interactome are either not annotated or have putative individualized functions that cannot be assigned to a broad category.

**Fig. 4.**
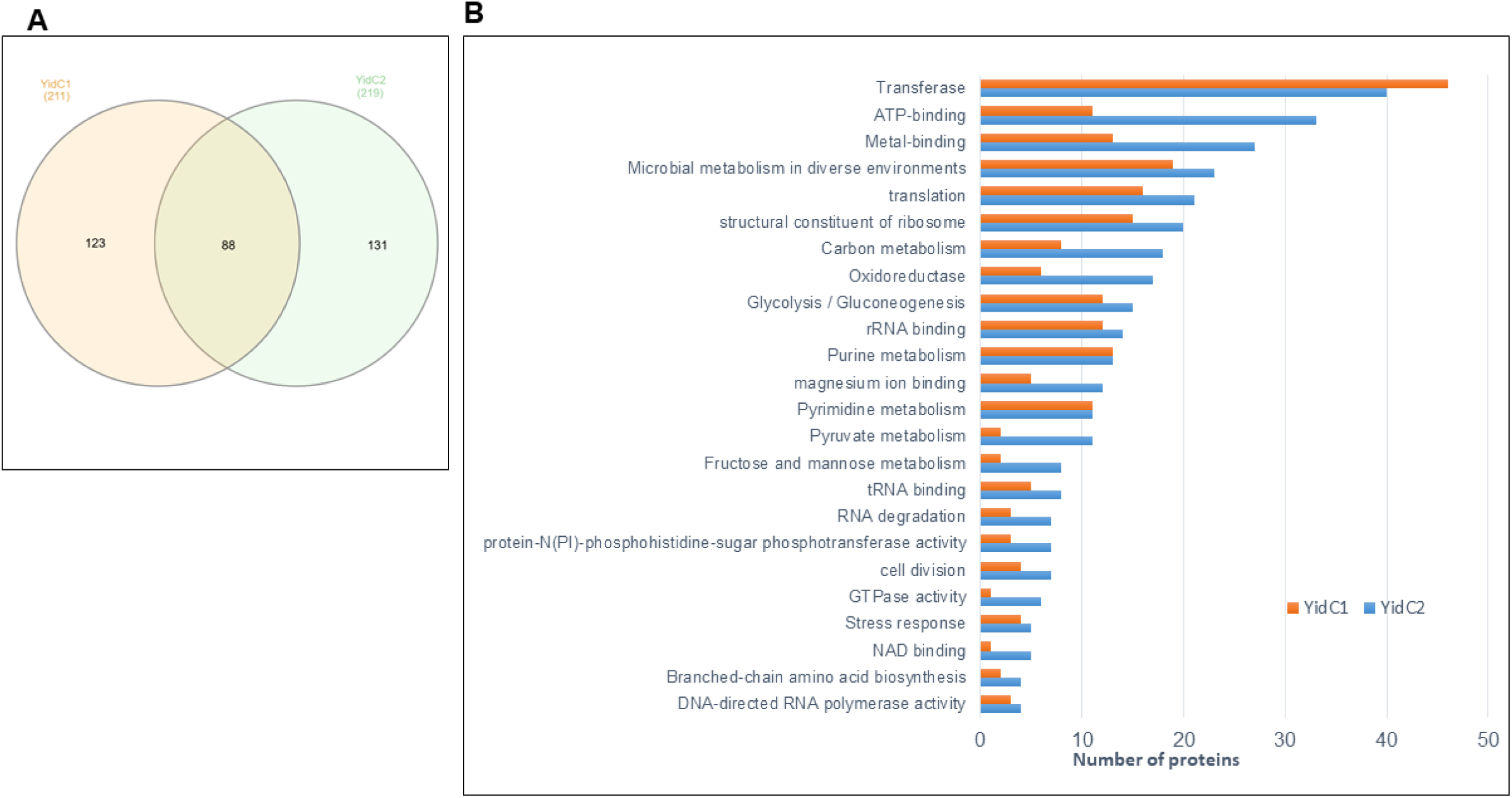
Comparison of YidC1 and YidC2 interactomes. **(A)** Venn diagram illustrating the degree of overlap of proteins identified in YidC1 and YidC2 interactomes. **(B)** Distribution of proteins within the YidC1 and YidC2 interactomes among various functional categories (DAVID analysis).

We also carried out a protein-protein interaction (PPI) network analysis using the STRING (Search Tool for the Retrieval of Interacting Genes/Proteins). YidC1 and YidC2, as well as all proteins experimentally identified as associating with either or both of them, were included in the uploaded datasets. The individual YidC1 and YidC2 STRING interactomes are shown in Fig. 5A and 5B, and the common interactome in Fig 5C. The majority of the proteins we identified in the current study were included within the PPI networks predicted by STRING, thus giving us high confidence in the accuracy of the experimentally determined protein interactomes. Consistent with co-translational protein translocation pathways, the most intense nodes identified in all three PPI network predictions were largely comprised of ribosomal proteins and other components of the translation machinery. L2, which we determined by ELISA not to interact with the YidC C-terminal tails (Fig. S3), was not predicted by STRING analysis to interact with either YidC1 or YidC2. S1 however, is a predicted STRING interaction partner of L2, as well as of YidC1 and YidC2. S1 is therefore a likely bridging molecule since it was detected experimentally whenever L2 was found in association with either YidC1 or YidC2.

**Fig. 5.**
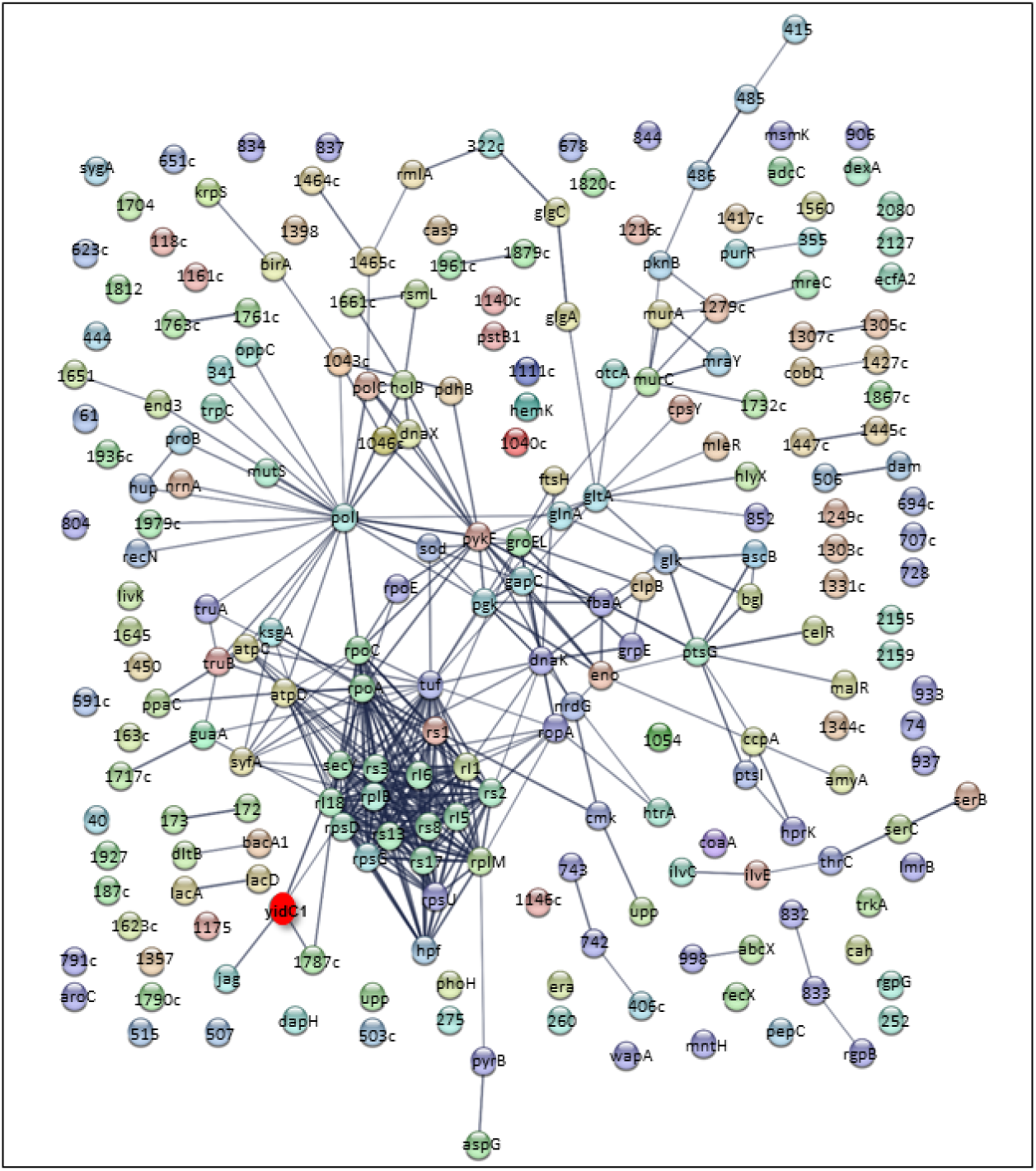
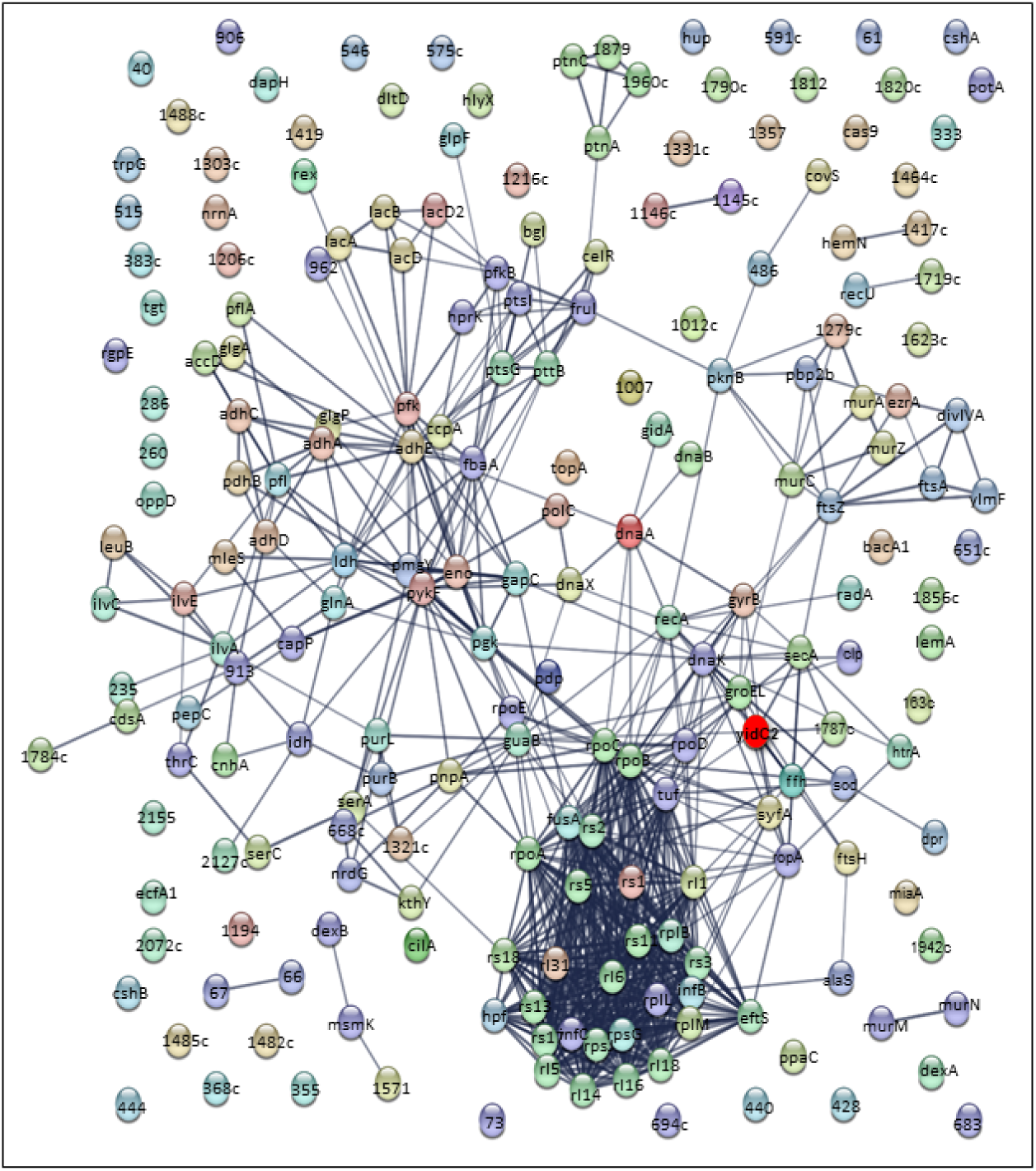
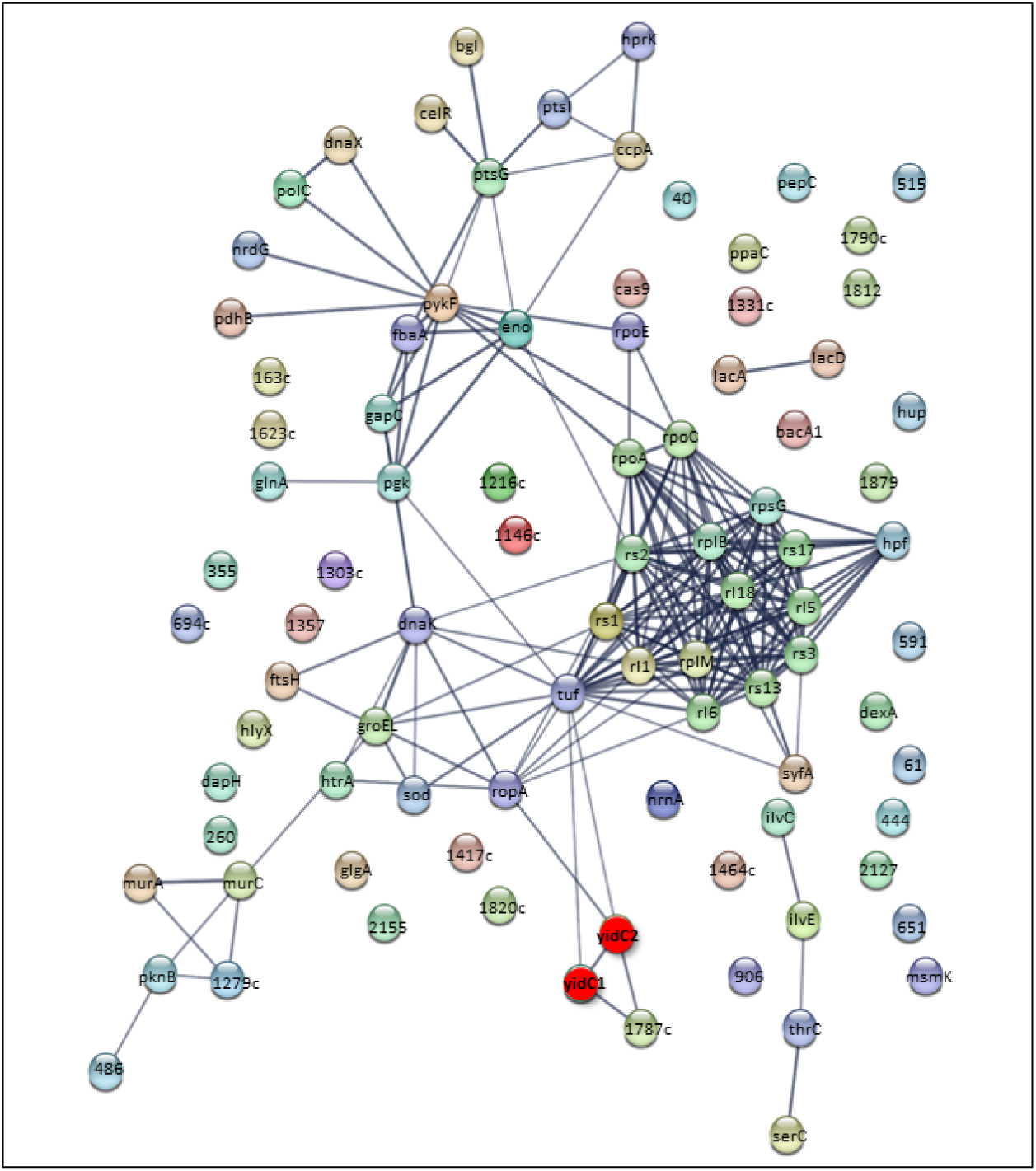
Protein-protein interaction networks predicted by STRING analysis. **(A)** YidC1 interactome **(B)** YidC2 interactome **(C)** YidC1 and YidC2 shared interactome. Each protein experimentally determined in the current study to associate with YidC1 and/or YidC2 is depicted by a sphere with either name or SMU number indicated. YidC1 and YidC2 are highlighted in red. Lines indicate predicted interactions based on current information within the STRING database.

### Concluding Remarks and Apparent *S. mutans* Protein Translocation Pathways

Most information regarding bacterial membrane protein translocation comes from the Gram-negative bacterium, *E. coli*; however, Gram-positive bacteria generally have two YidCs and, based on genomic sequences, a seemingly smaller holotranslocon whereby SecDF are lacking in streptococci and staphylococci. Our current results reveal that there are at least three putative pathways of membrane protein translocation in *S. mutans* : 1) SRP-YidC2, 2) SecY-YidC1, and 3) YidC1 and/or 2 only (Fig. 6). Ribosomal proteins were associated with all three apparent pathways consistent with the well accepted idea that insertion of membrane proteins is co-translational. In *S. mutans*, the majority of membrane protein substrates appear to prefer the SecY-YidC1 pathway since most of the predicted membrane proteins we identified were present in in the lower SDS-PAGE region in gel shift experiments where SecY and YidC1 were also found. A SecY-YidC interaction has been reported in *E. coli* under conditions of SecYEG overexpression [17]. Although our data does not yet prove a direct interaction between SecY and YidC1, their co-capture under endogenous conditions strongly suggests them to be a part of a common macromolecular complex. On a similar note, an SRP-YidC2 interaction was identified both by whole cell cross-linking gel shift, and GST-YidC2CT pull-down assays. Koch and coworkers also reported a cross-link between *E.coli* YidC and Ffh under endogenous conditions [17]. Our data are consistent with this result, and further demonstrate that there is a preference of YidC2 over YidC1 in working in concert with the SRP in the Gram-positive bacterium *S. mutans*. Eliminating YidC2 or SRP components apparently maim, but do not fully disable, this essential pathway. Hence YidC2 and SRP pathway deletion mutants are viable but stress sensitive, and double deletion of *yidC2* and *ffh* is lethal. Partitioning of a smaller number of putative substrates with the *S. mutans* SRP-YidC2 pathway components is consistent with the speculation that the SecY-YidC1 pathway is more likely the housekeeping mechanism for insertion of numerous membrane proteins under routine growth conditions, while the SRP-YidC2 pathway inserts membrane proteins necessary for survival of environmental stressors. We did not identify Ffh or FtsY as constituents of the YidC1/2 autonomous pathway; therefore, how such nascent substrate proteins are targeted to the membrane remains unclear. In mammalian cells, large ribosomal subunit proteins attach to the endoplasmic reticulum membrane to facilitate membrane targeting [60, 61]. We speculate therefore that a similar mechanism may exist in *S. mutans*. In support of this conjecture, L14, L16, and L31 were the only ribosomal proteins identified by gel shift assay in the middle region of the gel corresponding to the YidC1/2 autonomous pathway. Taken together, the current results add to our understanding of the organization and respective substrates of distinct protein transport pathways in a Gram-positive bacterium. This information will facilitate future research regarding the underlying biology of a prevalent oral pathogen.

**Fig. 6.**
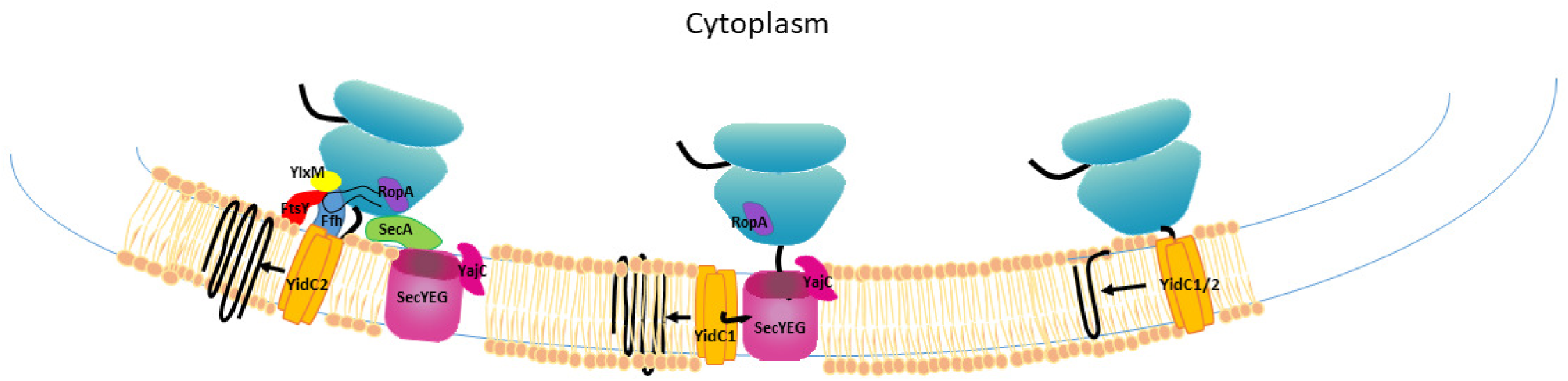
Model representation of putative co-translational membrane protein insertion pathways in *S. mutans*. (Left) SRP-YidC2 pathway. YidC2 works in concert with the signal recognition particle (SRP) pathway. The SRP is comprised of Ffh, a small cytoplasmic RNA, and the YlxM accessory protein present only in Gram-positive bacteria [63]. The SRP targets the ribosome nascent chain complex to the membrane via a reversible interaction of Ffh with the SRP receptor FtsY. The substrate protein is then passed to YidC2 for integration into the membrane. RopA and SecA fractionate with components of this pathway because of their common association with large ribosomal subunit proteins. **(Center) SecY-YidC1 pathway.** Integral membrane proteins are targeted to SecYEG with the help of RopA or other chaperones (DnaK, GroEL) and insertion into the membrane is facilitated by YidC1. **(Right) YidC1 and/or YidC2 autonomous pathway.** A small subset of membrane proteins with one or two transmembrane domains can be inserted into the membrane independently of SecYEG or the SRP.

## Materials and Methods

### Bacterial strains and media

*S. mutans* strains included NG8 [62], PC398 (*ΔyidC2* in NG8), and UA159. Strain PC398 was generated by PCR amplifying the allelic replacement cassette locus of strain AH398 (*ΔyidC2* in UA159) [7], transforming NG8 with the amplified DNA, selection on erythromycin, and sequence confirmation of the mutant construction. All cultures were grown at 37° C in Todd-Hewitt broth (BBL, Becton Dickinson) supplemented with 0.3%. yeast extract (THYE). Erythromycin (10 μg/ml) was added when appropriate. *E. coli* strain BL21 was grown aerobically at 37° C in Luria-Bertani (LB) broth or agar supplemented with ampicillin (100 μg/ml) or kanamycin (50 μg/ml) where appropriate.

### Formaldehyde cross-linking and Western blotting of whole cell lysates

Paraformalaldehyde (Sigma-Aldrich) was added to 4% (w/v) in phosphate buffered saline (PBS), pH 7.4, and stirred at 65° C with drop by drop addition of 1M NaOH until dissolution was complete. The solution was cooled to room temperature (RT), adjusted to pH 7.4, filtered (0.22 μm), and stored at 4°C for up to 4 weeks. Fifty ml of cells from mid-log phase *S. mutans* cultures (OD_600_ ∼ 0.6) were harvested by centrifugation at 5,300 x *g* for 30 min at 4°C, and washed twice with 10 ml PBS. The cell pellet was resuspended in 9.6 ml 0.4% formaldehyde solution and incubated for 15 min at 37°C with gentle shaking (Biometra OV5 3107A INC). The optimal concentration of 0.4% formaldehyde was established in pilot titration experiments. The reaction was quenched by addition of 0.4 ml 250 mM Tris, pH 7.4 (final concentration of 10 mM), and Sincubation at 37°C for 15 min. Paraformaldehyde-treated cells were pelleted by centrifugation and washed twice with PBS as above and resuspended to a final volume of 1 ml. Control cells were handled in the same way without formaldehyde. Whole-cell lysates were prepared from the cross-linked and untreated cell suspensions by glass bead breakage in a Mini-Bead Beater 8 apparatus (BioSpec Products, Inc., Bartlesville) for four 40 second cycles with 1 min cooling on ice between each cycle. Cell lysate samples were electrophoresed on 4-20% precast gels (Bio-Rad Laboratories, Hercules, CA) in Tris-Glycine-SDS buffer. Replicate gels were stained with Coomassie Blue R 250 or transblotted onto Immobilon polyvinylidene difluoride (PVDF) membranes (Bio-Rad Laboratories, Hercules, CA), reacted with affinity-purified YidC1 or YidC2 C-terminal-specific polyclonal rabbit antibodies (1:1000) [26], or anti-Ffh or anti-FtsY polyclonal rabbit antisera (1:1000) [63], followed by horseradish peroxidase-labeled anti-rabbit IgG (MP Biomedicals, Irvine, CA) (1:5000), and developed using the enhanced-chemiluminescence (ECL) Western blotting system (GE Healthcare).

### Coupling of anti-YidC2 antibodies to DynaBeads™ and immunocapture of protein complexes

Five mg of M-280 Tosylactivated Dynabeads™ (Invitrogen) were washed twice with 1 ml 0.1 M Na-phosphate buffer pH 7.4. The beads were coupled to affinity-purified rabbit polyclonal YidC2-specific antibodies generated against synthetic peptides corresponding to the C-terminal tail (NPPKPFKSNARKDITPQANNDKKLIT) and cytoplasmic loop 1 between TM2 and TM3 (SEKMAYLKPVFDPIQERMKNC). Beads were reacted at 37°C overnight with slow end over end rotation (Roto-Torque, Cole-Parmer, Chicago Illinois) in a final volume of 150 ul in 0.1 M Na-phosphate buffer pH 7.4 containing 50 ug of each purified antibody preparation and 3 M ammonium sulfate. Following incubation, the tube was placed next to a magnet and the supernatant removed. Unbound antibodies were removed from the beads by washing first with 1 ml PBS containing 0.5% TritonX-100, and secondly with freshly made 0.5 N NH_4_OH, 0.5 mM EDTA, until A_280_ of the wash supernatant was zero. Ab-coated beads were washed three times with 1 ml PBS, resuspended in 100 μl PBS, and reacted with ∼700 ul formaldehyde cross-linked whole cell lysate samples (∼ 8 mg/ml) derived from *S. mutans* strain NG8 (wild type) or PC398 (Δ*yidC2*), or with control samples prepared without formaldehyde, for three hours at 4° C with gentle end over end rotation. Next, beads were separated with a magnet and washed six times with 1 ml PBS. Ab-captured proteins were eluted with 0.5 ml freshly made 0.5 N NH_4_OH, 0.5 mM EDTA, and vortexing in an Eppendorf tube adapter (Vortex Mixer, Fisher Scientific) set at medium speed for 20 min at RT. Beads were removed with a magnet and the eluate was snap-frozen in liquid nitrogen and dried overnight at RT in a SpeedVac vacuum concentrator (Savant, Famingdale, NY). Twenty microliters of SDS sample buffer (62.5 mM Tris pH 6.8, 10 % Glycerol, 0.2 % SDS, 0.02% Bromphenol Blue) were added to each dried sample and incubated for 10 min at 65°C. The samples were electrophoresed on 4-20% gradient SDS-polyacrylamide gels (Bio-Rad, Hercules, CA) and analyzed by Western blot using anti-YidC2 C-terminal-specific antibodies as described above. Controls included non-cross-linked samples prepared without formaldehyde, and a Western blot developed with HRP-conjugated goat anti-rabbit secondary antibody only.

### Preparation of gel slices for protein identification by mass spectrometry

SDS polyacrylamide gels were rinsed in Optima LC-MS grade water (Fisher Scientific) three times, fixed for 15 min with 50% methanol and 7% acetic acid (Fisher Scientific), and stained with GelCode, Blue Stain Reagent (Thermo Scientific) according to the manufacturer’s instructions. Gel slices corresponding to gel-shifted regions identified by Western blot with anti-YidC1, YidC2, Ffh or FtsY-specific antibodies in the formaldehyde cross-linked UA159 whole cell lysate, but absent from the non-cross-linked control sample, were excised for *in situ* proteolysis. Similarly, a band detected by Western blot with anti-YidC2 antibodies in the DynaBead™ eluate of the NG8 formaldehyde cross-linked sample, but not the *ΔyidC2* mutant strain or non-cross-linked control samples, was excised for proteolysis from the same location of SDS-polyacrylamide gels of all four samples. Gel slices were washed twice in nanopure water for 5 minutes, then destained with 1:1 v/v methanol: 50 mM ammonium bicarbonate for ten minutes with two changes. Gel slices were dehydrated with 1:1 v/v acetonitrile: 50 mM ammonium bicarobonate, then rehydrated and incubated with dithiothreitol (DTT) solution (25 mM in 100 mM ammonium bicarbonate) for 30 minutes prior to the addition of 55 mM Iodoacetamide in 100 mM ammonium bicarbonate solution. Gel slices were incubated for an additional 30 min in the dark then washed with two cycles of water and dehydrated with 1:1 v/v acetonitrile: 50 mM ammonium bicarbonate. Protease was driven into the gel pieces by rehydrating them in 12 ng/ml trypsin in 0.01% ProteaseMAX Surfactant (Promega) for 5 minutes. Gel pieces were next overlaid with 40 µL of 0.01% ProteaseMAX surfactant: 50 mM ammonium bicarbonate and gently mixed on an orbital shaker for 1 hour. The digestion was stopped by addition of 0.5% trifluoroacetic acid. MS analysis was performed immediately to ensure high quality tryptic peptides with minimal non-specific cleavage.

### Mass spectrometry analysis

Nano-liquid chromatography tandem mass spectrometry (Nano-LC/MS/MS) was performed on a Thermo Scientific Q Exactive HF Orbitrap mass spectrometer equipped with an EASY Spray nanospray source (Thermo Scientific) operated in positive ion mode, or on a Quadrupole-Tof (Q-TOF) instrument. The LC system was an UltiMate™ 3000 RSLCnano system from Thermo Scientific. The mobile phase A was water containing 0.1% formic acid acetic acid and the mobile phase B was acetonitrile with 0.1% formic acid. Five microliters of each sample was first injected on to a Thermo Fisher Scientific Acclaim Trap Cartridge (C18 column, 75 um ID, 2 cm length, 3 μm 100 Å pore size) and washed with mobile phase A to desalt and concentrate the peptides. The injector port was switched to inject and the peptides were eluted off of the trap onto the column. An EASY Spray PepMAP column from Thermo Scientific was used for chromatographic separations (C18, 75 um ID, 25 cm length, 3 μm 100 Å pore size). The column temperature was maintained 35° C as peptides were eluted directly off the column into the LTQ system using a gradient of 2-80%B over 45 minutes, with a flow rate of 300 nL/min. The total run time was 60 minutes. The MS/MS was acquired according to standard conditions established in the lab. The EASY Spray source operated with a spray voltage of 1.5 KV and a capillary temperature of 200° C. The scan sequence of the mass spectrometer was based on the TopTen™ method; the analysis was programmed for a full scan recorded between 350 – 2000 Da, and a MS/MS scan to generate product ion spectra to determine amino acid sequence in consecutive instrument scans of the ten most abundant peak in the spectrum. The AGC Target ion number was set at 30,000 ions for full scan and 10,000 ions for MSn mode. Maximum ion injection time was set at 20 ms for full scan and 300 ms for MSn mode. Micro scan number was set at 1 for both full scan and MSn scan. The CID fragmentation energy was set to 35%. Dynamic exclusion was enabled with a repeat count of 1 within 10 seconds, a mass list size of 200, and an exclusion duration 350 seconds. The low mass width was 0.5 and the high mass width was 1.5.

### Database searching

All MS/MS samples were analyzed using Sequest (XCorr Only) (Thermo Fisher Scientific, San Jose, CA, USA; version IseNode in Proteome Discoverer 2.2.0.388 or Mascot Server 2.7). Sequest (XCorr Only) was set up to search *Streptococcus mutans* UA159 or NG8 (GenBank: AE014133.2 GenBank: CP013237.1, respectively). Sequest (XCorr Only) was searched with a fragment ion mass tolerance of 0.020 Da and a parent ion tolerance of 10.0 PPM.

### Criteria for protein identification

Scaffold (version Scaffold_4.8.6, Proteome Software Inc., Portland, OR) was used to validate MS/MS based peptide and protein identifications. Peptide identifications were accepted if they could be established at greater than 95.0% probability by the Peptide Prophet algorithm [64] with Scaffold delta-mass correction. Protein identifications were accepted if they could be established at greater than 99.0% probability and contained at least 1 identified peptide. Protein probabilities were assigned by the Protein Prophet algorithm [65].

### Two-dimensional difference gel electrophoresis (DIGE) analysis of *S. mutans* proteins captured with GST-YidC1 compared to GST-YidC2 C-terminal tail fusion proteins

The C-terminal fragment (bp682-816) of *yidC1* was amplified by PCR using primers NL5F (ggaacggatcccaggtcttccagattctgttg) and NL5R (ccgtagtcgacttattttctctttttatgtgctttc). The C-terminal fragment (bp742-933) of *yidC2* was amplified by PCR using primers NL6F (ggaacggatccacaaaccatatcattaaaccaaaat) and NL6Rb (ccgtagtcgacttattgcttatggtgacgctgt). *S. mutans* UA159 genomic DNA was used as the template. PCR products were digested with *BamH*I and *Sal*I and ligated to corresponding restriction enzyme sites in the pGEX-4T-2 vector. The vector only encoding GST was transformed into BL21 DE3 (ThermoFisher Scientific). Plasmids encoding GST-YidC1CT or GST-YidC2CT were transformed into BL21 Star™ (ThermoFisher). GST-YidC1CT expression was induced with 0.5 mM isopropyl-ß-D-thiogalactopyranoside (IPTG) for 6 hours at 30° C. Expression of GST and GST-YidC2CT was induced with 1 mM IPTG for 4 hours at 37° C. Cells were harvested by centrifugation at 11,325 x *g* for 15 min, and resuspended in 25 ml PBS. Cell suspensions were supplemented with 1 mM phenylmethylsulfonyl fluoride (PMSF) (Acros Organics) and protease inhibitor cocktail (1 mini tablet/25 ml) (Roche Diagnostics GmbH). Cell lysis was performed using an Avestin EmulsiFlex-C5 high-pressure homogenizer (Avestin Inc., Ottawa, Ontario, Canada) at a pressure of 15,000-20,000 p.s.i. for three cycles. Cell debris was removed by centrifugation at 11,000 x *g* for 30 min and the supernatants filtered through a 0.22 µm syringe filter (Merck Millipore). Recombinant proteins were purified on an AKTA Purifier system (GE Healthcare) using a GSTrap column and elution with 50 mM Tris-HCl, 10 mM reduced glutathione pH 8.0. Purified proteins were dialyzed in equilibration buffer (50 mM Tris, 150 mM NaCl, pH 8.0) and incubated with Pierce® Glutathione Spin Columns (Thermo Scientific) at RT for 1 hour with gentle rotation according to the manufacturer’s instructions. A fresh whole cell lysate of *S. mutans* UA159 was filtered through a 0.22 µm syringe filter (Merck Millipore) and incubated with the GST and GST fusion proteins bound Glutathione Spin Columns overnight at 4°C with gentle rotation. The column was washed four times with PBS and bound proteins were eluted with 50 mM Tris, 150 mM NaCl, pH8 containing 10 mM reduced glutathione. Eluates were separated by electrophoresis through 4-20% gradient SDS-polyacrylamide gels (Bio-Rad, Hercules, CA) and visualized by Coomassie blue staining to confirm protein capture, then sent to Applied Biomics (Hayward, CA) on dry ice for proteomic analysis. Proteins captured with GST, GST-YidC1CT, or GST-YidC2CT were labeled with CyDye DIGE blue Cy5, red Cy3, or green cy2 fluors respectively, separated on a single 2D gel electrophoresis and the gel was analyzed for spot picking, followed by trypsin digestion for Mass Spectrometry protein identification. Peptides were subjected to tandem matrix-assisted laser desorption ionization-time of flight (MALDI-TOF) for peptide mass fingerprinting and MALDI-TOF/TOF for identification of peptide sequences which were searched against *S. mutans* UA159 database from NCBI and SwissProt using MASCOT search engine (Matrix Science). Proteins with Protein Score or Total Ion, confidence interval (C.I.) greater than 95% were considered significant.

### Bioinformatic analyses

Amino acid sequences of proteins identified in all the experiments were downloaded from the *S. mutans* strain NG8 assembly database (https://www.ncbi.nlm.nih.gov/nuccore/CP013237.1) or UA159 assembly database (https://www.ncbi.nlm.nih.gov/nuccore/AE014133.2), and analyzed for the presence and number of transmembrane domains using the webtool, TMHMM v2.0 [66]. Functional analysis of proteins was conducted using the Database for Annotation, Visualization, and Integrated Discovery (DAVID) bioinformatics tool 6.8 (http://david.abcc.ncifcrf.gov/) [67]. Protein-protein interaction (PPI) network analysis was performed using the STRING (Search Tool for the Retrieval of Interacting Genes/Proteins) database with a minimum required interaction score set to high confidence (0.700) [68]. YidC1 and YidC2 were manually added to the respective analyses as these proteins themselves were not the part of uploaded datasets.

### Expression and purification of recombinant proteins

Recombinant *E. coli* were induced to produce 50S ribosomal protein L2 or trL2 with 0.05 mM IPTG at RT overnight. Bacterial cells were harvested by centrifugation at 11,325 x *g* for 15 min, and resuspended in 25 ml 50 mM sodium phosphate, 300 mM sodium chloride,10 mM imidazole, pH 7.4, supplemented with 1 mM phenylmethylsulfonyl fluoride (PMSF) (Acros Organics) and protease inhibitor cocktail (1 mini tablet/25 ml) (Roche Diagnostics GmbH). Cell lysis was performed using an Avestin EmulsiFlex-C5 high-pressure homogenizer (Avestin Inc., Ottawa, Ontario, Canada) at a pressure of 15,000-20,000 p.s.i. for three cycles. Cell debris was removed by centrifugation and the supernatant was filtered through a 0.22 µm syringe filter (Merck Millipore). Recombinant proteins were purified on an AKTA Purifier system (GE Healthcare) using a HiTrap TALON column and eluted with 50 mM sodium phosphate, 300 mM sodium chloride, Ph. 7.4, containing 150 mM imidazole.

## Acknowledgements

This work was supported by NIH NIDCR award DE008007 to LJB and funding from NIH S10 OD021758 to the University of Florida Mass Spectrometry Research and Education Center. We thank Dr. Kari B. Basso and Dr. Manasi Kamat of the Department of Chemistry, University of Florida for technical expertise and mass spectrometry sample analysis.

